# Accuracy of non-parametric species richness estimators across taxa and regions

**DOI:** 10.1101/2022.08.23.504921

**Authors:** Arttu Soukainen, Pedro Cardoso

## Abstract

1. Non-parametric species richness estimators are efficient and widely used when sampling is incomplete. There is little consensus on which of the available estimators works best across taxa and regions. Until now no work compared existing algorithms with multiple datasets encompassing contrasting scenarios.
2. We used data from 62 inventories worldwide at different spatial scales, including 20 vertebrate, 22 invertebrate and 20 plant datasets, and compared the accuracy of the most used non-parametric estimators (Chao and Jackknife) and improvements to their original formulations.
3. Our results highlight the good performance of the Jackknife estimators for incidence data, especially the P-corrected first order jackknife estimator (Jack1inP). This algorithm ranked most often the best or among the best performing estimators using two measures of accuracy that measure deviation from expectation along the accumulation curve.
4. We argue that Jack1inP can be considered a universal estimator for species richness, regardless of taxon, temporal and spatial scales, or completeness of the sampling. More research should however be directed towards finding the precise contexts when each estimator might perform best.

## Introduction

Biodiversity is a multidimensional concept, but species richness remains the most reported facet. Although conceptually easy to quantify, species richness is not always easy to measure. On a global scale, we are nowhere near answering the question of how many species there are. More than 2.1 million eukaryotic species have been described to date (IUCN 2021), but it is estimated that 86% of terrestrial and 91% of marine species are still unknown to science (Mora et al. 2011). We do have a reasonably accurate picture of some terrestrial groups, such as vertebrates, vascular plants, and a few groups of insects such as butterflies, dragonflies, and bees. This is due to the invaluable work of taxonomists over the last centuries. Still, for other groups like most insects, arachnids, nematodes, fungi and microorganisms, the taxonomic and biogeographic knowledge remains limited (Cardoso et al. 2011). Even when we look at more limited areas and focus on a specific taxon, measuring species richness can prove challenging, especially in the case of hyper-diverse groups.

Local species richness is ideally determined through extensive sampling, where each organism is individually identified. Sampling is continued until new species no longer appear and the species accumulation curve is complete. In most cases, however, this can be difficult or practically impossible (Basset et al. 2012). Research resources are often insufficient, and sampling is carried out with variable levels of effort. For example, the study area may simply be too large, or the sampling methods inadequate to detect all species. As a result, observed species richness almost invariably underestimates the true richness. A significant problem is created by hyper-diverse taxa such as invertebrates in tropical regions. Despite ambitious attempts, the identification of all species, even from relatively small areas of rainforest, have fallen short (Basset et al. 2012). In those cases, assessing true richness based on the observed richness can be highly misleading. Obtaining accurate estimates of species richness from incomplete sampling therefore seems essential.

Because there are undetected species in most taxonomic surveys or species inventories, researchers use extrapolation methods to predict the true number of species. Several extrapolation methods have been proposed and discussed in the literature, and some of these are commonly used in ecological research (Bunge and Fitzpatrick 1993; Chao and Chiu 2016; Colwell and Coddington 1994). There are three main approaches to species richness estimation: 1) parametric estimators based on the species-abundance curves; 2) fitting asymptotic formulas to randomized accumulation curves; and 3) nonparametric estimators.

Parametric estimators are based on the concept of ‘veil-line’, assuming that species abundances are log-normally distributed in natural communities and deviations are due to undersampling (Colwell and Coddington 1994). Species accumulation curves are fitted with asymptotic formulas whose asymptote represents the true diversity of a site (Soberon and Llorente 1993). Non-parametric approaches aim to control the dependence of the observed richness on sample completeness and sampling effort. Non-parametric estimators do not assume a specific form of the species abundance distribution. Instead, to estimate the number of undetected species, these approaches use information on the frequency of observed rare species in a sample (Chao and Chiu 2016).

Most tests to date found that non-parametric estimators are the best at predicting true richness across multiple settings (Walther and Moore 2005), probably because they rely on less assumptions about the structure of communities (O’Hara 2005). Although non-parametric estimators have been found to outperform other estimator classes, it is unclear which of the specific estimators works best among the dozen or so proposed. Previous works almost invariably tested estimators on single or very few datasets or with limited geographical, temporal and taxonomic extents (Poulin 1998; Brose 2002; Chao and Chiu 2016; Ter Steege et al. 2017; Brito et al. 2021; Hortal et al. 2006). It is not surprising then, that previous tests are conflicted in their conclusions. No work previously tested a range of estimators under contrasting scenarios covering a wide range of variables that can influence estimator performance.

Here we compare the most commonly used non-parametric estimators, including some advancements that were seldom tested after being proposed (Lopez et al. 2012). These estimators can be divided into abundance and incidence-based depending on the input data required. We use several different datasets representing different taxa, regions, spatial and temporal extents for comparison. Finally, we assess whether there are clear differences between the different estimators when comparing their accuracy values across contrasting datasets.

## Methods

Field sampling data was collected from a variety of open online sources. We searched for articles where species richness estimation was used, and data from taxonomic inventories or long-term surveys. In the article search, we used Google Scholar search for (species OR communit*) AND (inventor* OR estimat*). Online sources included datasets from GBIF (Global Biodiversity Information Facility: https://www.gbif.org/dataset/search), Data Dryad (https://datadryad.org/search) and the LTER network (Long Term Ecological Research: https://portal.edirepository.org). The searches were conducted between September 2020 and October 2021. The last search was performed on October 7, 2021. We sought to cover different regions, taxonomic groups, and spatial and temporal extents without the need to be exhaustive, but with the explicit goal of being comprehensive in the coverage of these dimensions. In addition, we had strict criteria for datasets to qualify, as we needed to evaluate the accuracy of estimators against reasonable targets:

1. The dataset was collected using standardized sampling methods and sampling was done in such a way that it was possible to divide the data in samples, so that incidence-based methods could be tested.
2. The dataset contained spatial and temporal metadata so that both space and time could be quantified and compared.
3. The observed species richness had to be 10 at minimum, so that using estimators can be seen as useful.
4. The dataset had to be complete or close to complete, i.e. the sampling was continued until new species no longer appeared. As fully complete data sets were difficult to obtain, the main criteria were sampling effort (number of individuals divided by number of species) above 50 or sampling effort above 25 and number of singletons (species with a single individual in the dataset) less than 20%. These criteria increased the probability that most estimators were able to reach unbiased (no systematic deviation from true values), precise (small variance) and congruent (small differences between estimators) values when the complete datasets were used.

With the datasets we collated metadata regarding taxon, region, spatial extent, and temporal extent. For all datasets we calculated the species richness, number of samples, total abundance, number of singletons, doubletons (species with two individuals in the dataset), uniques (number of species occurring in only one sample), and duplicates (number of species occurring in two samples). If there was no clear way to determine the spatial extent, it was marked as zero, for example, cases of highly mobile species like birds that were sampled with point observations or flying insects that were sampled with single traps. For each dataset, it was important to determine what was considered as a sample. If no clear definition was provided with the data, this was done by us according to some natural way to divide the entire data. For example, individuals collected from a particular plot on a given day were considered as one sample. The following nonparametric estimators were calculated.

First order jackknife estimator for abundance data (Jack1ab) (Burnham and Overton 1978):

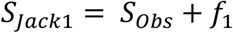

Where *S*_*Obs*_ is the observed number of species (species richness) in the sample and *f*_1_ is the number of singletons in the sample. First order jackknife estimator for incidence data (Jack1in):

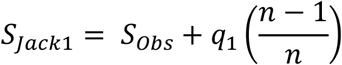

Where *q*_1_ is the number of uniques and *n* is the number of samples. Second order jackknife estimator for abundance data (Jack2ab) (Smith and van Belle 1984):

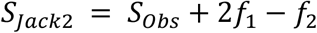

Where *f*_2_ is the number of doubletons. Second order jackknife estimator for incidence data (Jack2in):

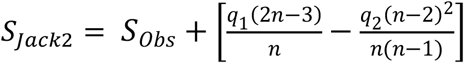

Where *q*_2_ is the number of duplicates. Chao 1 estimator for abundance data (Chao 1984)

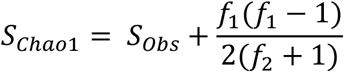

And Chao 2 estimator for incidence data (Chao 1987):

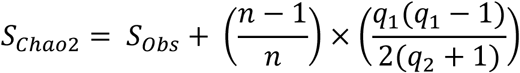

The P-corrected version of each of the six previous estimators was also calculated (Lopez et al. 2012):

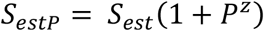

Where *S*_*est*_ is the original estimate, *P* is the proportion of singletons/uniques, and *z* is calculated as:

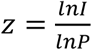

Where *lnI* is the natural logarithm of the sampling effort and *lnP* is the natural logarithm of the proportion of singletons/uniques.

We then compared the accuracy values of the estimators. The accuracy of the estimator depends on its ability to predict the target value. In order to account for the possibility of incomplete datasets, we used the average of all estimators as the target. In this case the accuracy was thus defined as the overall distance between the estimated values and the target value (Walther and Moore 2005) along the rarefaction curve. The unweighted accuracy values were calculated as:

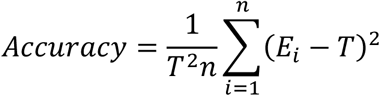

Where *T* is the average of the estimators as target value and *E*_*i*_ is the estimator value for the i:th sample. In addition, the weighted accuracy values were calculated as:

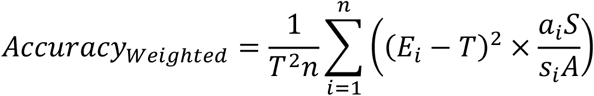

Where *a*_*i*_ is the abundance (number of individuals) in the i:th sample, *A* is the total abundance, *s*_*i*_ is the number of species in the i:th sample and *S* is the total species richness. Weighted accuracy gives higher weight to the samples at the end of the accumulation curve, benefiting estimators that do not diverge from true richness when sampling is high, meaning that estimated values are allowed to largely fluctuate with few samples. Unweighted accuracy gives the same weight to any point in the curve, meaning that estimators should be accurate also for low sampling effort, but with the caveat that they might diverge with high effort, which is uncommon but plausible. None of these versions can be considered *a priori* preferable.

In addition to the 12 estimators described above, the accuracy of the observed species rarefaction curve was also calculated, as a minimal benchmark that any estimator worth considering must outperform. For calculating the estimated values and respective accuracy, we used the *alpha*.*accum* function from the R package *BAT* (Cardoso et al. 2015). For each dataset, estimators and the observed richness were given a rank from 1 to 13 (1 being the most accurate). To provide an overall picture of the ranking data, marginal and pairwise frequencies were calculated with the *pmr* package (Lee and Yu 2013) in R. In addition, we calculated weighted mean rank and variance, where the same weight was given to each source of data, i.e., paper, to account for multiple datasets being taken from single papers, this way avoiding overweighting datasets with similar characteristics. We also used the Wilcoxon signed-rank test (Wilcoxon 1945), to compare the estimator with lowest mean rank with every other estimator.

## Results

From 49 different sources, we collected a total of 62 datasets for comparison, including 20 representing vertebrates, 22 invertebrates and 20 flora *sensu lato* (plants and fungi). North America, South America, Africa, Oceania, Europe and Asia were represented with 13, 4, 9, 6, 22, and 8 datasets, respectively. The spatial extent varied from 0.5ha to 1700 km2. The temporal extent from four days to 36 years. For more information on the datasets see Appendix A. Calculated species richness estimates for each dataset are presented in Appendix B. Marginal frequencies for all estimators are presented in Appendix C.

According to the weighted mean, P-corrected Jack1 incidence estimator (Jack1inP), had the best overall rank, for both weighted and unweighted accuracy (Table I). Testing the hypothesis that Jack1inP has better rank than any other estimator in the comparison (*H*_0_: *Rank*_*Jack*1*inP*_ ≥ *Rank*_*E*_, *H*_1:_ *Rank*_*Jack*1*inP*_ *< Rank*_*E*_). The Wilcoxon signed rank test indicated that the ranks of Jack1inP were statistically significantly better than ranks of all other estimators, in both unweighted and weighted accuracy comparisons. Jack1inP ranked first 37 times out of 62 and was ranked in the top three in 90% of datasets for unweighted accuracy and in 80% for weighted accuracy (Figs. 1 and 2). As expected, observed species richness was the least accurate overall.

**Table I.** Mean of rank and standard deviation of accuracy (unweighted and weighted) of all estimators in order of lowest mean rank across 62 datasets. All comparisons with the best performing estimator (Jack1inP) are significant (Wilcoxon p-value < 0.001 in all cases).

| Unweighted |  |  | Weighted |  |  |
| --- | --- | --- | --- | --- | --- |
| Estimator | Mean rank | SD | Estimator | Mean rank | SD |
| Jack1inP | 1.85 | 1.22 | Jack1inP | 2.63 | 2.46 |
| Jack2in | 3.67 | 2.18 | Jack1in | 3.80 | 2.39 |
| Jack1in | 5.07 | 2.41 | Jack2in | 4.25 | 2.99 |
| Jack2inP | 5.13 | 4.16 | Jack1abP | 5.30 | 2.13 |
| Jack2abP | 5.17 | 2.33 | Jack2abP | 5.81 | 2.43 |
| Jack1abP | 5.91 | 2.12 | Jack2ab | 6.26 | 2.04 |
| Jack2ab | 6.34 | 1.85 | Jack1ab | 6.92 | 3.06 |
| Jack1ab | 7.59 | 2.64 | Jack2inP | 7.03 | 3.88 |
| Chao2 | 8.80 | 1.96 | Chao2 | 8.63 | 3.27 |
| Chao1P | 8.80 | 3.17 | Chao1P | 8.66 | 2.04 |
| Chao1 | 10.14 | 1.90 | Chao1 | 9.89 | 2.50 |
| Chao2P | 11.08 | 3.02 | Chao2P | 10.50 | 3.07 |
| Observed | 11.44 | 2.05 | Observed | 11.33 | 2.57 |

**Fig. 1.**
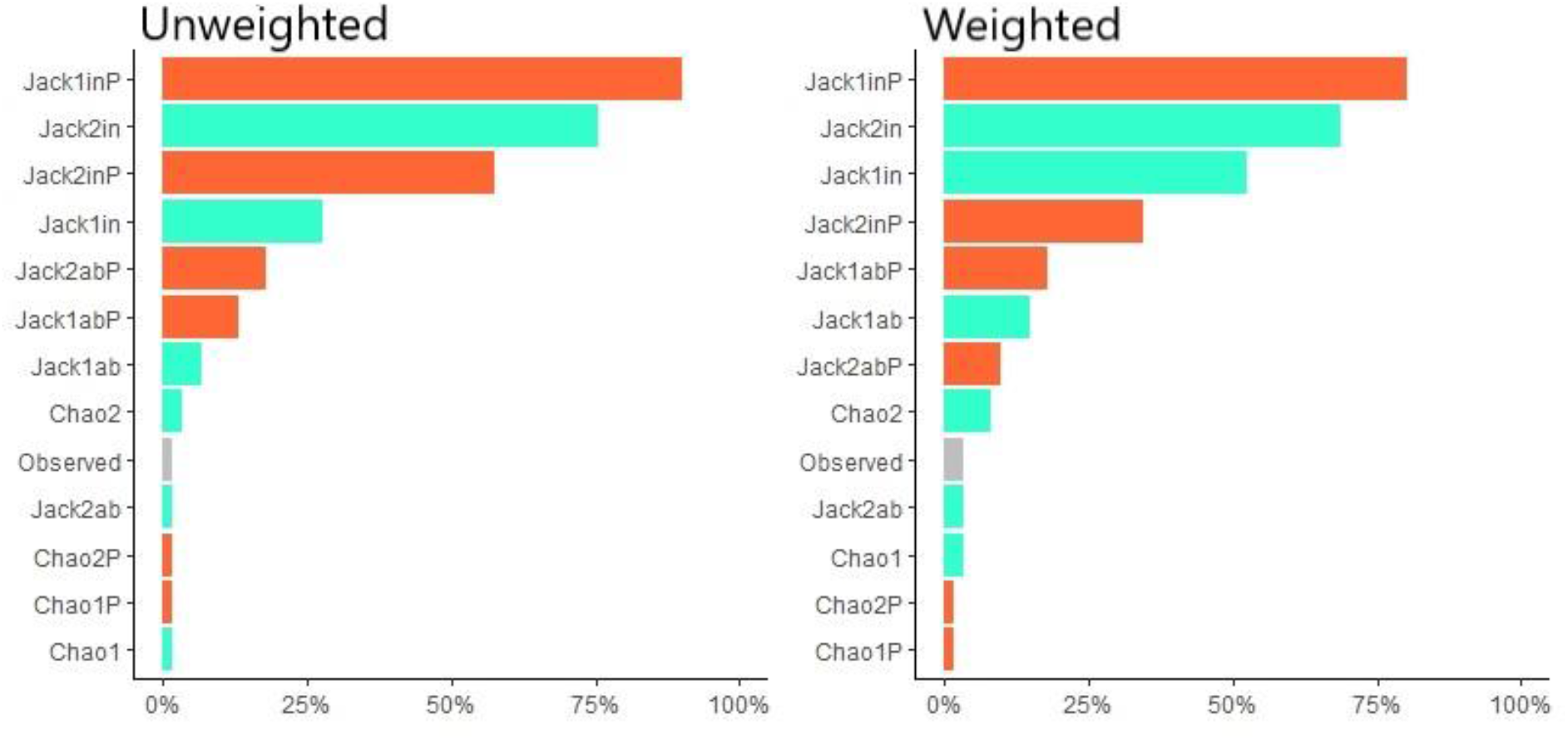
Percentage of how many times each estimator was ranked in the top three according to unweighted or weighted accuracy.

**Fig. 2.**
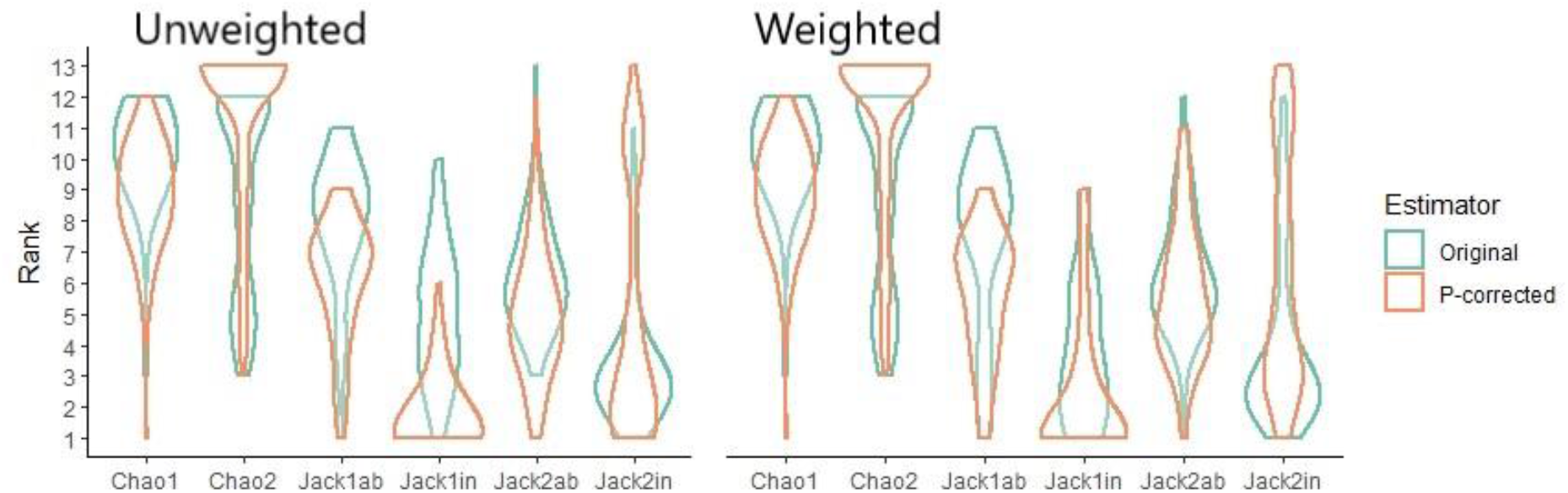
Rank distribution of estimator accuracy across 62 datasets.

For unweighted accuracy, in addition to Jack1inP, Jack2in and Jack2inP ranked in the top three on over 50% of the datasets. In comparison, Jack1inP ranked higher than Jack2in and Jack2inP, 53 and 45 times (out of 62) respectively. Pairwise frequencies for all estimators are presented in Appendix D. The ranking distributions of the estimators reflect the overall results, with jack1inP being the best estimator across all datasets (Fig. 2).

For weighted accuracy: Jack1inP, Jack1in and Jack2in ranked in the top three for over 50% of the datasets. Jack1inP ranked higher than Jack1in and Jack2in, 45 and 52 times respectively. The ranking distributions of the estimators reflect the overall results, with jack1inP being the best estimator across all datasets (Fig. 2).

## Discussion

Our results highlight the P-corrected first order Jackknife estimator for incidence data (Jack1inP) to be the best performing estimator in this comparison of a broad range of datasets. These results are consistent for both unweighted accuracy, which considers the sampling process, and weighted accuracy, which benefits estimators which fall closer to the target value at the end of the sampling. Jack1inP has by far the best overall rank in both comparisons, suggesting that Jack1inP is the best estimator for any level of completeness of the datasets, stabilizing its values around the target value with relatively few samples but also in cases of more complete sampling.

It should be noted that because not all the datasets used were complete, the target value was set to the average of all estimators at the end of the sampling, not the true species richness. In case of close to complete datasets, this target value can be considered a reasonably accurate prediction of the actual species richness. In any case, both weighted and unweighted accuracies consider the full sampling process, not only the final value, hence using the average will not benefit estimators that fall close to it with full datasets. Lopez et al. (2012) proposed the P-correction for reducing estimator bias in cases of undersampling. They suggested that datasets with less than 8% of singletons or sampling effort less than 100 probably have some degree of undersampling. Of our 62 datasets, 20 failed to meet this threshold which could explain the better ranking for P-corrected estimators in these cases, however, the success of Jack1inP extends beyond this, almost always ranking as the best or among the best performing estimators. We present a strong case that P-correction not only improves the performance of Jack1in in cases of undersampling, but overall for variable levels of completeness. The descriptive statistics provide some indication that P-correction improved the overall ranking of some of the other estimators as well, but not as clearly as with Jack1inP. For example, the original Jack2in estimator had more top 3 rankings than the P-corrected version.

The four best performing estimators for both unweighted and weighted accuracy, Jack1inP, Jack2in, Jack2inP and Jack1in are all estimators for incidence data. Incidence data lacks information on the number of individuals and only indicates whether the species is present or absent in the samples. For example, if we have sampling units spread across an area, incidence data provides a list of species in each of these samples. These estimators have the advantage that they only require sample data and can be used with datasets without abundance information. All the datasets we collected contained abundance data, but based on our results, abundance information is not always paramount for obtaining a reasonably accurate estimate. Collecting incidence data may require less resources in some cases, and for certain groups of organisms, incidence data can be the most practical way of presenting data. For example, when dealing with organisms like corals or mosses, it may not make sense to assess the number of individuals, but to determine which species are present on a particular plot. Newer methods using next generation sequencing for determining species richness may not provide abundance data (Hiiesalu et al. 2012; Lindeque et al. 2013), often only resulting in a species list per sample. In those cases, using nonparametric incidence-based estimators is the only viable option. On the other hand, incidence-based estimators present the disadvantage that they require data to be divided in samples, and the way samples are divided might influence the results (Hortal et al. 2006).

In case of unweighted accuracy, Jack2inP was the estimator to most often outperform Jack1inP (Appendix D), but it also has numerous worst ranks, making it the estimator with most variance within the ranks. Looking at the weighted accuracy, the overall ranking of Jack2inP drops out of the best performing estimators. This is probably a result of overestimation in several cases, however, it is out of the scope of this paper to determine the common factor for cases where jack2inP is the most efficient estimator and where it might overestimate. Overall, the jackknife estimators performed better in this comparison, than the Chao estimators. Some of the previous comparisons of nonparametric estimators in the literature have reached similar conclusions (Chiarucci et al. 2003; Branco et al. 2018). In our results, the original Chao2 was the best Chao estimator.

Many of the previous comparisons of nonparametric estimation methods in the literature have been limited to a particular taxonomic group, with varying results. Basualdo (2011) suggested that the best estimators for benthic macroinvertebrates would be Jack1, Chao1 and ACE (Abundance-based Coverage Estimator). Conversely, Chiarucci et al. (2003) concluded that Jack2 would be the best estimator in a high diversity plant community, although none of the estimators appeared accurate enough. Brose (2002) in turn, suggested that for carabid beetles caught by pitfall traps, the Chao 2 would be the most accurate. The number of datasets used in our comparison is not sufficient to draw conclusions if and how the accuracy of estimators change with taxon. Differences in the performance of estimators do not necessarily depend on the taxon but rather on the environment. Distribution of species abundances in space, e.g., alpine plants, is likely to follow different rules than their relatives in the rainforest. The completeness of the samples may also affect which estimator works best and this may well depend on the taxon under consideration. Acquiring complete datasets with abundance data is often close to impossible, except for some sessile groups of organisms such as trees. For example, with flying insects such as Lepidoptera, as a result of extensive sampling, unique species are always expected, and the number of unique species is likely to grow steadily in long term surveys. It is not uncommon for an insect to travel long distances because of a weather phenomenon, for example, which then causes a single point observation (Rainey 1989).

In our literature search we found no previous reports of P-corrected Jack1in as the most accurate nonparametric estimator. Using extensive datasets covering a large taxonomic scope and contrasting experimental setups, Jack1inP turned out to be the best choice for estimating species richness in most settings, by far. Should there be evidence of a certain estimator being most accurate in particular settings, choosing that method might be advisable. Otherwise, Jack1inP is likely to provide a reasonably accurate prediction of species richness, regardless of sampling completeness, taxon, or sampling arrangement. We therefore recommend using this estimator as a universal algorithm independently of the many variables that might influence the accuracy of the different formulas proposed to date.

## References

Basset, Y.L., et al. (2012) Arthropod diversity in a tropical forest. Science, 338: 1481. https://doi.org/10.1126/science.1226727.

Basualdo, C.V. (2011) Choosing the best non-parametric richness estimator for benthic macroinvertebrates databases. Revista de la Sociedad Entomológica Argentina, 70: 27–38.

Branco, M., Francisco G.F., Cermeño, P. (2018) Assessing the efficiency of non-parametric estimators of species richness for marine microplankton. Journal of Plankton Research, 40: 230–243. https://doi.org/10.1093/plankt/fby005.

Brito, P.G., Jovem-Azevêdo, D., Campos, M.A., Paiva, F.F., Molozzi, J. (2021) Performance of richness estimators for invertebrate inventories in reservoirs. Environmental Monitoring and Assessment, 193: 686. https://doi.org/10.1007/s10661-021-09487-z

Brose, U. (2002) Estimating species richness of pitfall catches by non-parametric estimators. Pedobiologia, 46: 101–107. https://doi.org/10.1078/0031-4056-00117.

Bunge, J., Fitzpatrick, M. (1993) Estimating the number of species: a review. Journal of the American Statistical Association, 88: 364–373. https://doi.org/10.1080/01621459.1993.10594330.

Burnham, K.P., Overton, W.S. (1978) Estimation of the size of a closed population when capture probabilities vary among animals. Biometrika, 65: 625–633. https://doi.org/10.1093/biomet/65.3.625.

Cardoso, P., Erwin, T.L., Borges, P.A.V., New, T.R. (2011) The seven impediments in invertebrate conservation and how to overcome them. Biological Conservation, 144: 2647–2655. https://doi.org/10.1016/j.biocon.2011.07.024

Cardoso, P., Rigal, F. & Carvalho, J.C. (2015) BAT - Biodiversity Assessment Tools, an R package for the measurement and estimation of alpha and beta taxon, phylogenetic and functional diversity. Methods in Ecology and Evolution, 6: 232–236. https://doi.org/10.1111/2041-210X.12310

Chao, A. (1984) Nonparametric estimation of the number of classes in a population. Scandinavian Journal of Statistics, 11: 265–270.

Chao, A. (1987) Estimating the population size for capture-recapture data with unequal catchability. Biometrics, 43: 783–791. https://doi.org/10.2307/2531532.

Chao, A., Chiu, C.H. (2016) Nonparametric estimation and comparison of species richness. eLS. https://doi.org/10.1002/9780470015902.a0026329.

Chiarucci, A., Enright, N.J., Perry, G.L.W., Miller, B.P., Lamont, B.B. (2003) Performance of nonparametric species richness estimators in a high diversity plant community. Diversity and Distributions, 9: 283–295. https://doi.org/10.1046/j.1472-4642.2003.00027.x.

Colwell R.K, Coddington J.A. (1994) Estimating terrestrial biodiversity through extrapolation. Philosophical Transactions of the Royal Society of London. Series B: Biological Sciences, 345: 1311. https://doi.org/10.1098/rstb.1994.0091.

Hiiesalu, I., Opik, M., Metsis, M., Lilje, L., Davison, D., Vasar, M., Moora, M., Zobel, M., Wilson, S.D., Pärtel, M. (2012) Plant species richness belowground: higher richness and new patterns revealed by next-generation sequencing. Molecular Ecology, 21: 2004–2016. https://doi.org/10.1111/j.1365-294X.2011.05390.x

Hortal, J., Borges, P.A.V., Gaspar, C. (2006) Evaluating the performance of species richness estimators: sensitivity to sample grain size. Journal of Animal Ecology, 75: 274–287. https://doi.org/10.1111/j.1365-2656.2006.01048.x

International Union for Conservation of Nature (2021) Red List Summary Statistics. Available online at: https://www.iucnredlist.org/resources/summary-statistics. Accessed 09.08.2022.

Lee, P.H., Yu, P.L.H. (2013) An R package for analyzing and modeling ranking data. BMC Medical Research Methodology, 13: 65. https://doi.org/10.1186/1471-2288-13-65.

Lindeque, P.K., Parry, H.E., Harmer, R.A., Somerfield, P.J., Atkinson, A. (2013) Next generation sequencing reveals the hidden diversity of zooplankton assemblages. PloS One, 8: e81327. https://doi.org/10.1371/journal.pone.0081327

Lopez, L.C.S., Fracasso, M.P.A., Mesquita, D.O., Palma, A.R.T., Riul, P. (2012) The relationship between percentage of singletons and sampling effort: a new approach to reduce the bias of richness estimates. Ecological Indicators, 14: 164–169. https://doi.org/10.1016/j.ecolind.2011.07.012.

Mora, C., Tittensor, D.P., Adl, S., Simpson, A.G.B., Worm, B. (2011) How many species are there on Earth and in the ocean? PLoS Biology 9: e1001127. https://doi.org/10.1371/journal.pbio.1001127

O’Hara, R.B. (2005) Species richness estimators: how many species can dance on the head of a pin? Journal of Animal Ecology, 74: 375–386. https://doi.org/10.1111/j.1365-2656.2005.00940.x

Poulin, R. (1998) Comparison of three estimators of species richness in parasite component communities. Journal of Parasitology, 84: 485–490. https://doi.org/10.2307/3284710

Rainey, R.C. (1989) Migration and Meteorology: Flight Behaviour and the Atmospheric Environment of Locusts and Other Migrant Pests. Oxford University Press, USA.

Smith, E.P., van Belle, G. (1984) Nonparametric estimation of species richness. Biometrics, 40: 119–129. https://doi.org/10.2307/2530750.

Soberon J., Llorente, J. (1993) The use of species accumulation functions for the prediction of species richness. Conservation Biology, 7: 480–488.

Ter Steege, H., Sabatier, D., Oliveira, S.M., Magnusson, W.E., Molino, J.-F., Gomes, V.F., Pos, E.T., Salomão, R.P. (2017) Estimating species richness in hyper-diverse large tree communities. Ecology, 98: 1444–1454. https://10.1002/ecy.1813

Walther, B.A., Moore, J.L. (2005) The concepts of bias, precision and accuracy, and their use in testing the performance of species richness estimators, with a literature review of estimator performance. Ecography, 98: 1444–1454. https://doi.org/10.1111/j.2005.0906-7590.04112.x.

Wilcoxon, F. (1945) Individual comparisons by ranking methods. Biometrics Bulletin, 1: 80–83. https://doi.org/10.2307/3001968.

