## Appendix C for "Accuracy of non-parametric species richness estimators across taxa and regions"

**Table C1.** Marginal frequencies for unweighted accuracy ranks. Number of times an estimator was ranked at a particular rank among 62 datasets. Estimators in order of lowest mean rank.

| | 1 | 2 | 3 | 4 | 5 | 6 | 7 | 8 | 9 | 10 | 11 | 12 | 13 |
|---|---|---|---|---|---|---|---|---|---|---|---|---|---|
| **Jack1inP** | 37 | 14 | 4 | 4 | 1 | 1 | 0 | 0 | 0 | 0 | 0 | 0 | 0 |
| **Jack2in** | 2 | 20 | 24 | 5 | 3 | 0 | 3 | 1 | 2 | 0 | 1 | 0 | 0 |
| **Jack1in** | 2 | 6 | 9 | 12 | 5 | 13 | 5 | 5 | 1 | 3 | 0 | 0 | 0 |
| **Jack2inP** | 14 | 11 | 10 | 2 | 6 | 0 | 1 | 2 | 0 | 6 | 6 | 2 | 1 |
| **Jack2abP** | 1 | 6 | 4 | 16 | 12 | 10 | 4 | 4 | 2 | 1 | 0 | 1 | 0 |
| **Jack1abP** | 1 | 4 | 3 | 3 | 8 | 6 | 25 | 6 | 5 | 0 | 0 | 0 | 0 |
| **Jack2ab** | 0 | 0 | 1 | 5 | 16 | 18 | 7 | 9 | 2 | 2 | 0 | 0 | 1 |
| **Jack1ab** | 3 | 0 | 1 | 4 | 1 | 2 | 8 | 12 | 18 | 6 | 6 | 0 | 0 |
| **Chao2** | 0 | 0 | 2 | 6 | 7 | 4 | 1 | 3 | 6 | 2 | 7 | 23 | 0 |
| **Chao1P** | 1 | 0 | 0 | 0 | 1 | 4 | 4 | 16 | 11 | 15 | 8 | 1 | 0 |
| **Chao1** | 0 | 0 | 1 | 1 | 0 | 1 | 1 | 2 | 11 | 17 | 15 | 13 | 0 |
| **Chao2P** | 0 | 0 | 1 | 1 | 1 | 3 | 2 | 0 | 3 | 3 | 0 | 5 | 42 |
| **Observed** | 0 | 0 | 1 | 1 | 0 | 0 | 0 | 1 | 0 | 6 | 18 | 16 | 18 |

**Table C2.** Marginal frequencies for weighted accuracy ranks. Number of times an estimator was ranked at a particular rank among 62 datasets. Estimators in order of lowest mean rank.

| | 1 | 2 | 3 | 4 | 5 | 6 | 7 | 8 | 9 | 10 | 11 | 12 | 13 |
|---|---|---|---|---|---|---|---|---|---|---|---|---|---|
| **Jack1inP** | 37 | 7 | 5 | 3 | 1 | 3 | 0 | 3 | 2 | 0 | 0 | 0 | 0 |
| **Jack1in** | 10 | 12 | 10 | 8 | 4 | 11 | 0 | 3 | 3 | 0 | 0 | 0 | 0 |
| **Jack2in** | 2 | 24 | 16 | 5 | 2 | 2 | 0 | 4 | 1 | 3 | 0 | 2 | 0 |
| **Jack1abP** | 3 | 2 | 6 | 6 | 9 | 7 | 21 | 4 | 3 | 0 | 0 | 0 | 0 |
| **Jack2abP** | 1 | 2 | 3 | 17 | 11 | 7 | 10 | 1 | 5 | 1 | 3 | 0 | 0 |
| **Jack2ab** | 1 | 1 | 0 | 4 | 17 | 17 | 6 | 6 | 5 | 3 | 0 | 1 | 0 |
| **Jack1ab** | 3 | 4 | 2 | 4 | 2 | 1 | 6 | 14 | 12 | 9 | 4 | 0 | 0 |
| **Jack2inP** | 3 | 5 | 13 | 5 | 6 | 2 | 4 | 4 | 2 | 3 | 5 | 5 | 4 |
| **Chao2** | 0 | 2 | 3 | 7 | 3 | 2 | 0 | 5 | 6 | 9 | 6 | 18 | 0 |
| **Chao1P** | 0 | 0 | 1 | 1 | 2 | 2 | 9 | 11 | 13 | 11 | 9 | 2 | 0 |
| **Chao1** | 1 | 1 | 0 | 1 | 0 | 1 | 1 | 5 | 6 | 15 | 13 | 17 | 0 |
| **Chao2P** | 0 | 1 | 0 | 0 | 3 | 4 | 4 | 1 | 2 | 5 | 5 | 4 | 32 |
| **Observed** | 0 | 0 | 2 | 0 | 1 | 2 | 0 | 0 | 1 | 2 | 16 | 12 | 25 |
