## Appendix D for "Accuracy of non-parametric species richness estimators across taxa and regions"

**Table D1.** Pairwise frequencies for unweighted accuracy ranks. Number of times a given estimator was ranked higher than each of the other estimators. Total of 62 data sets in the comparison.

| | Jack1inP | Jack2in | Jack1in | Jack2inP | Jack2abP | Jack1abP | Jack2ab | Jack1ab | Chao1P | Chao2 | Chao1 | Chao2P | Observed |
|---|---|---|---|---|---|---|---|---|---|---|---|---|---|
| **Jack1inP** | - | 53 | 58 | 45 | 57 | 58 | 59 | 58 | 60 | 61 | 60 | 60 | 60 |
| **Jack2in** | 8 | - | 47 | 36 | 48 | 52 | 53 | 52 | 57 | 58 | 57 | 60 | 58 |
| **Jack1** | 3 | 14 | - | 25 | 33 | 43 | 37 | 52 | 52 | 54 | 58 | 55 | 61 |
| **Jack2inP** | 16 | 25 | 36 | - | 42 | 42 | 43 | 43 | 47 | 53 | 46 | 60 | 50 |
| **Jack2abP** | 4 | 13 | 28 | 19 | - | 45 | 53 | 49 | 57 | 46 | 57 | 56 | 57 |
| **Jack1abP** | 3 | 9 | 18 | 19 | 16 | - | 24 | 58 | 57 | 43 | 61 | 52 | 59 |
| **Jack2ab** | 2 | 8 | 24 | 18 | 8 | 37 | - | 47 | 56 | 43 | 57 | 53 | 57 |
| **Jack1ab** | 3 | 9 | 9 | 18 | 12 | 3 | 14 | - | 41 | 36 | 57 | 51 | 61 |
| **Chao1P** | 1 | 4 | 9 | 14 | 4 | 4 | 5 | 20 | - | 32 | 58 | 48 | 59 |
| **Chao2** | 0 | 3 | 7 | 8 | 15 | 18 | 18 | 25 | 29 | - | 31 | 54 | 38 |
| **Chao1** | 1 | 4 | 3 | 15 | 4 | 0 | 4 | 4 | 3 | 30 | - | 46 | 59 |
| **Chao2P** | 1 | 1 | 6 | 1 | 5 | 9 | 8 | 10 | 13 | 7 | 15 | - | 19 |
| **Observed** | 1 | 3 | 0 | 11 | 4 | 2 | 4 | 0 | 2 | 23 | 2 | 42 | - |

**Table D2.** Pairwise frequencies for weighted accuracy ranks; Number of times a given estimator was ranked higher than each of the other estimators. Total of 62 data sets in the comparison.

| | Jack1inP | Jack1in | Jack2in | Jack1abP | Jack2abP | Jack2ab | Jack1ab | Jack2inP | Chao2 | Chao1P | Chao1 | Chao2P | Observed |
|---|---|---|---|---|---|---|---|---|---|---|---|---|---|
| **Jack1inP** | - | 45 | 52 | 50 | 56 | 53 | 50 | 56 | 58 | 57 | 56 | 61 | 56 |
| **Jack1in** | 16 | - | 27 | 49 | 42 | 44 | 54 | 38 | 59 | 54 | 59 | 57 | 61 |
| **Jack2in** | 9 | 34 | - | 46 | 50 | 48 | 47 | 53 | 53 | 52 | 54 | 57 | 55 |
| **Jack1abP** | 11 | 12 | 15 | - | 27 | 31 | 54 | 27 | 47 | 58 | 57 | 53 | 59 |
| **Jack2abP** | 5 | 19 | 11 | 34 | - | 46 | 45 | 29 | 43 | 53 | 53 | 55 | 55 |
| **Jack2ab** | 8 | 17 | 13 | 30 | 15 | - | 39 | 28 | 44 | 54 | 54 | 52 | 56 |
| **Jack1ab** | 11 | 7 | 14 | 7 | 16 | 22 | - | 22 | 40 | 42 | 58 | 48 | 61 |
| **Jack2inP** | 5 | 23 | 8 | 34 | 32 | 33 | 39 | - | 43 | 44 | 44 | 50 | 49 |
| **Chao2** | 3 | 2 | 8 | 14 | 18 | 17 | 21 | 18 | - | 29 | 35 | 52 | 42 |
| **Chao1P** | 4 | 7 | 9 | 3 | 8 | 7 | 19 | 17 | 32 | - | 55 | 44 | 58 |
| **Chao1** | 5 | 2 | 7 | 4 | 8 | 7 | 3 | 17 | 26 | 6 | - | 39 | 58 |
| **Chao2P** | 0 | 4 | 4 | 8 | 6 | 9 | 13 | 11 | 9 | 17 | 22 | - | 26 |
| **Observed** | 5 | 0 | 6 | 2 | 6 | 5 | 0 | 12 | 19 | 3 | 3 | 35 | - |
