## Appendix A for "Accuracy of non-parametric species richness estimators across taxa and regions"

**Appendix A.** Metadata for all datasets used in this work.

| Name | Reference | Taxon | Region | Spatial extent (km2) | Dates | Temporal extent (days) | Sample definition | Number of samples (q) | Total abundance (n) | Number of species (S) | Sampling effort (n/S) | % singletons |
|---|---|---|---|---|---|---|---|---|---|---|---|---|
| **amphibiansTua** | Costa et al. 2021 | Amphibia | River Tua, Portugal | 250 | November 15, 2010 - May 26, 2017 | 2384 | Direct observations at sample point | 336 | 2844 | 12 | 237 | 8.33 |
| **annelidaBelarus** | Baturina et al. 2021 | Annelida | Naroch lakes, Belarus | 79.6 | January 1, 1997 - December 31, 2018 | 8034 | single lake bottom sample | 155 | 5220 | 52 | 100 | 21.15 |
| **batsKruger** | Parker 2019 | Chiroptera | Kruger National Park, South Africa | 30 | January 1, 2017 - December 31, 2017 | 365 | Observations from specific location | 14 | 574 | 23 | 25 | 4.35 |
| **beesBurkina1** | Coulibaly and Stein 2018 | Bees | Burkina Faso | 0 | September 30, 2013 - October 30, 2015 | 760 | Sampling event | 113 | 21435 | 36 | 595 | 0 |
| **beesBurkina2** | Coulibaly and Stein 2018 | Bees | Burkina Faso | 0 | September 30, 2013 - October 30, 2015 | 760 | Sampling event | 102 | 14924 | 37 | 403 | 2.7 |
| **beesBurkina3** | Coulibaly and Stein 2018 | Bees | Burkina Faso | 0 | September 30, 2013 - October 30, 2015 | 760 | Sampling event | 88 | 2449 | 37 | 66 | 0 |
| **beesCanada** | Ratti 2004 | Bees | Fraser Valley, British Columbia, Canada | 0 | April 15, 2003 - July 1 2004 | 443 | observations from single day | 91 | 6175 | 63 | 98 | 12.7 |

| | | | | | | | | | | | | |
|---|---|---|---|---|---|---|---|---|---|---|---|---|
| **beetlesRussia** | Makarov and Matalin 2020 | Ground beetles | Elton lake, Russia | 100 | May 10, 2006 - May 10, 2007 | 365 | Sampling event (Specimens from "sampling line") | 207 | 51314 | 148 | 347 | 13.51 |
| **benthicMed1** | Chatzinikolaou and Arvanitidis 2017 | Benthos (Annelida, Mollusca, Arthropoda, Echinodermata, Nemertea, Sipuncula | El Kantaoui, Tunisia | 0 | February 13, 2012 - September 25, 2012 | 225 | single sample from a specific station | 45 | 13938 | 172 | 81 | 18.6 |
| **benthicMed2** | Chatzinikolaou and Arvanitidis 2017 | Benthos (Annelida, Mollusca, Arthropoda, Echinodermata, Nemertea, Sipuncula | Heraklion, Crete, Greece | 0 | February 13, 2012 - September 25, 2012 | 225 | single sample from a specific station | 42 | 8611 | 131 | 66 | 22.9 |
| **benthicMed3** | Chatzinikolaou and Arvanitidis 2017 | Benthos (Annelida, Mollusca, Arthropoda, Echinodermata, Nemertea, Sipuncula) | Cagliari, Sardinia, Italy | 0 | February 13, 2012 - September 25, 2012 | 225 | single sample from a specific station | 75 | 15150 | 140 | 108 | 17.86 |
| **birdsAzores** | Goulart et al. 2019 | Aves | Terceira Island, Azores, Portugal | 0 | August 21, 2013 - October 31, 2018 | 1897 | Sampling event (15min bird observations) | 2003 | 82985 | 107 | 776 | 0 |
| **birdsBelgium** | Girotto and Groom 2018 | Aves | Meise Botanical Garden, Belgium | 0 | January 16, 2017 - June 20, 2017 | 18 | Birds observen from a transect | 36 | 1376 | 37 | 37 | 8.11 |
| **birdsKorea** | Choi et al. 2020 | Aves | Haseong-Myeon, South Korea | 0 | 2018-11-27 - 2019-4-26 | 150 | birds observed in single sampling period | 19 | 41697 | 112 | 372 | 13.39 |
| **chiroUkraine** | Bitušík et al. 2020 | Chironomidae | Carpathians, Ukraine | - | June 23, 2019 - June 26, 2019 | 4 | specimens from single pond | 8 | 1236 | 35 | 35 | 14.29 |
| **culicidaeBelgium** | Dekoninck et al. 2021 | Culicidae | Maasmechelen, Belgium | 0 | July 28, 2009 - November 20, 2009 | 115 | sampling event | 28 | 529 | 17 | 31 | 11.76 |
| **echinoMontenegro** | Kascelan et al. 2015 | Echinodermata | South Adriatic, Montenegro | - | January 1, 2005 - January 1, 2008 | 1095 | specimens collected by certain method on a single day | 36 | 8202 | 50 | 164 | 4 |

| | | | | | | | | | | | | |
|---|---|---|---|---|---|---|---|---|---|---|---|---|
| **epiphytesNicaragua** | Berrios and Hazel 2020 | Epiphytes | Volcan Maderas, Nicaragua | 1 | 09/07/2003 - July 14, 2003 | 6 | Epiphyte species identified from single tree | 40 | 17427 | 80 | 218 | 5 |
| **fishBotswana1** | Mosie and Makati 2020 | Fish | Okavango, Botswana | - | June 12, 2003 - October 27, 2016 | 4886 | fish captured with a net after 12 h | 99 | 19076 | 50 | 382 | 8 |
| **fishBotswana2** | Mosie and Makati 2020 | Fish | Okavango, Botswana | - | June 12, 2003 - October 27, 2017 | 4886 | fish captured with a net after 12 h | 23 | 44126 | 26 | 1697 | 11.54 |
| **fishBotswana3** | Mosie and Makati 2020 | Fish | Okavango, Botswana | - | June 12, 2003 - October 27, 2018 | 4886 | fish captured with a net after 12 h | 33 | 11064 | 32 | 346 | 9.38 |
| **fishCanada** | Cornthwaite 2021 | Fish | South Hard Bottom Longline, British Colombia, Canada | - | January 1, 2005 - December 31, 2018 | 5112 | a random depth-stratified design and the sampling units are 2 km by 2 km blocks | 36 | 1376 | 37 | 37 | 8.11 |
| **fishMexico** | Paz-Ríos et al. 2021 | Actinopteri, Elasmobranchii | Terminos Lagoon, Mexico | 1700 | January 1, 1980 - August 17, 2017 | 13712 | Sampling event period | 32 | 48717 | 141 | 346 | 18.44 |
| **fishMyanmar** | Kano et al. 2016 | Fish | Inle Lake, Myanmar | 120 | September 23, 2014 - July 2, 2016 | 648 | specimens from a certain habitat in and around the lake | 44 | 1362 | 41 | 33 | 4.88 |
| **fishSabor** | Beja et al. 2021 | Fish | River Sabor, Portugal | 3170 | June 14, 2012 - July 7, 2020 | 2945 | Sampling event | 302 | 27514 | 14 | 1965 | 7.14 |
| **fungiAzores** | Melo et al. 2020 | Glomeromycota | Terceira and São Miguel Islands, Azores, Portugal | - | August 14, 2007 - September 17, 2013 | 2226 | sampling event | 226 | 18747 | 37 | 507 | 2.7 |
| **lepidopteraSpain** | Barea-Azcón 2021 | Lepidoptera | Sierra Nevada, Spain | 0 | July 7, 2008 - March 8, 2020 | 4262 | walk transects from a single day | 1615 | 97905 | 102 | 960 | 0.98 |

| | | | | | | | | | | | | |
|---|---|---|---|---|---|---|---|---|---|---|---|---|
| **LTER_ants** | Ellison and Gotelli 2018 | Formicidae | Cornwall, NY, US | 0.1 | July 7, 2007 - July 1, 2015 | 2916 | Specimens collected from a plot on a given day | 52 | 4773 | 41 | 116 | 17.07 |
| **LTER_birds** | Bateman et al. 2018 | Aves | Phoenix metropolitan area, US | 0 | January 9, 2013 - February 5, 2018 | 1853 | Bird observation event | 189 | 34950 | 187 | 187 | 10.7 |
| **LTER_butterflies** | Ross 2014 | Rhopalocera | Oregon, US | 120 | July 6, 1994 - May 3, 1995 | 301 | butterflies collected on a single day | 84 | 3090 | 68 | 45 | 13.24 |
| **LTER_Orthoptera** | Haarstad 2018 | Orthoptera | Minnesota, US | 30 | May 30, 1992 - September 30 ,1992 | 123 | Specimen collected from certain plot | 49 | 9734 | 52 | 187 | 15.38 |
| **LTER_plants** | Seastedt 2019 | Plantae | Colorado, US | 5.2 | 2000 | - | plants from single "tree island" | 14 | 2337 | 43 | 54 | 11.63 |
| **LTER_reefFish1** | Brooks 2019 | Fish | Moorea, French Polynesia | 2.5 | July 11, 2006 - August 5, 2018 | 4408 | Specimens collected from a plot on a given day | 28 | 65593 | 266 | 247 | 13.91 |
| **LTER_reefFish2** | Brooks 2019 | Fish | Moorea, French Polynesia | 2.7 | July 11, 2006 - August 5, 2018 | 4408 | Specimens collected from a plot on a given day | 28 | 63506 | 260 | 244 | 13.46 |
| **LTER_reefFish3** | Brooks 2019 | Fish | Moorea, French Polynesia | 4 | July 11, 2006 - August 5, 2018 | 4408 | Specimens collected from a plot on a given day | 29 | 54322 | 257 | 211 | 14.4 |
| **LTER_reefFish4** | Brooks 2019 | Fish | Moorea, French Polynesia | 4.5 | July 11, 2006 - August 5, 2018 | 4408 | Specimens collected from a plot on a given day | 26 | 49709 | 256 | 194 | 12.89 |
| **LTER_reefFish5** | Brooks 2019 | Fish | Moorea, French Polynesia | 10 | July 11, 2006 - August 5, 2018 | 4408 | Specimens collected from a plot on a given day | 29 | 48721 | 251 | 194 | 11.95 |

| | | | | | | | | | | | | |
|---|---|---|---|---|---|---|---|---|---|---|---|---|
| **LTER_reefFish6** | Brooks 2019 | Fish | Moorea, French Polynesia | 5.8 | July 11, 2006 - August 5, 2018 | 4408 | Specimens collected from a plot on a given day | 27 | 59689 | 250 | 239 | 12 |
| **mammalsColombia** | Herrera 2020 | Large mammals | Montes de Maria, Colombia | - | May 12, 2018 - September 30, 2018 | 141 | sampling event | 62 | 769 | 25 | 31 | 4 |
| **molluscsCanaries** | Langeraert and Brosens 2020 | Snails | Gran Canaria, Spain | 1560 | August 3, 2016 - February 8, 2020 | 1284 | Specimens from specific location | 35 | 2904 | 54 | 54 | 3.7 |
| **odonataTua** | Costa et al. 2020 | Odonata | River Tua, Portugal | 0 | June 6, 2010 - August 4, 2017 | 2616 | 15 min observations from sampling point | 218 | 1802 | 40 | 45 | 10 |
| **orthopteraCastroVerde** | Pina 2017 | Orthoptera | Castro Verde, Portugal | 0 | April 17, 2014 - July 17, 2015 | 45 | Observations with same location ID | 320 | 2083 | 35 | 60 | 11.43 |
| **plantsIndonesia** | Schrader et al. 2020 | Woody plants | Gam Bay, Indonesia | - | June 2016 - February 2018 | 610 | Trees from one island | 40 | 2215 | 57 | 39 | 7.02 |
| **plantsLebanon** | El Zein 2020 | Vascular plants | Danniye-Akkar, Lebanon | 29 | June 1, 2018 - December 1, 2019 | 548 | single sampling quandrat | 160 | 28149 | 458 | 61 | 4.59 |
| **plantsRussia** | Trubina and Nesterkov 2021 | Vascular plants | Southern Urals, Russia | - | July 4, 2003 - July 14, 2003 | 10 | plants inside single sampling plot | 700 | 5585 | 73 | 77 | 12.33 |
| **plantsSpain** | Vega-Álvarez et al. 2019 | Plantae | Salamanca, Spain | 1.5 | May 2007 - June 2015 | 2953 | Plot/year | 539 | 43842 | 146 | 300 | 6.85 |
| **plantsSweden1** | Lind and Nilsson 2015 | Vascular plants | Västerbotten, Sweden | 0 | 2011 | ? | Plant coverage inside a sampling plot | 192 | 8979 | 86 | 104 | 16.28 |
| **plantsSweden2** | Lind and Nilsson 2015 | Vascular plants | Västerbotten, Sweden | 0 | 2012 | ? | Plant coverage inside a sampling plot | 189 | 8118 | 81 | 100 | 4.94 |

| | | | | | | | | | | | | |
|---|---|---|---|---|---|---|---|---|---|---|---|---|
| **plantsSweden3** | Lind and Nilsson 2015 | Vascular plants | Västerbotten, Sweden | 0 | 2013 | ? | Plant coverage inside a sampling plot | 165 | 14498 | 92 | 158 | 6.52 |
| **polinatorsBelgium** | Noel et al. 2021 | Anthophila + Syrphidae | Wallonia, Belgium | 0 | April 1, 2018 - July 31, 2019 | 486 | Specimens from specific habitat | 9 | 6220 | 120 | 52 | 20.83 |
| **spidersArrabida** | Cardoso et al. 2008a | Araneae | Arrabida Nature Park, Portugal | 0.01 | June 1 - 15, 2004 | 14 | Specimens captured by certain method from the plot | 320 | 5548 | 150 | 37 | 17.33 |
| **spidersGeres** | Cardoso et al. 2008b | Araneae | Peneda Geres Nature Park, Portugal | 0.01 | June 1 -15, 2005 | 14 | Specimens captured by certain method from the plot | 320 | 7516 | 186 | 40 | 19.89 |
| **treesBCI** | Pyke et al. 2001 | Trees | Barro Colorado Island, Panama | 5 | 1982 | - | Trees inside a single plot | 50 | 21457 | 225 | 95 | 8.44 |
| **treesBenin** | Mensah et al. 2020 | Trees | Benin | 0.223 | 2018? | ? | Trees inside a single plot | 40 | 8894 | 46 | 193 | 4.35 |
| **treesBrazil** | Gastauer 2016 | Trees | Fundo gallery forest, Bom Despacho region, Minas Gerais, Brazil | 0.005 | June 26, 2007 - August 2, 2015 | 2790 | Trees inside a plot | 51 | 1735 | 80 | 22 | 18.75 |
| **treesChina1** | Zhang et al. 2016 | Plantae | Mengsong, China | 100 | May, 2009 - September, 2011 | 882 | Plants from a single plot | 28 | 77775 | 807 | 96 | 16.6 |
| **treesChina2** | Zhang et al. 2016 | Trees | Yinggeling, China | 500 | May, 2009 - September, 2011 | 882 | Trees from a single plot | 29 | 10114 | 371 | 27 | 15.09 |
| **treesCocoli** | Condit 1998 | Trees | Cocoli river, Panama | 0.04 | December 1, 1998 - December 23, 1998 | 23 | Trees observed from the plot on a given day | 30 | 9425 | 175 | 54 | 17.14 |
| **treesPuertoRico** | Zimmerman et al. 2010 | Trees | Luquillo Forest, Puerto Rico | 0.16 | 2005 | ? | Trees inside a sampling quadrat | 400 | 115758 | 146 | 793 | 9.59 |
| **treesRwanda** | Nyirambangutse et al. 2017 | Trees | Nyungwe National Park, Rwanda | 0.075 | 2011 - 2012 | - | Trees inside a plot | 15 | 6161 | 86 | 72 | 9.3 |

| | | | | | | | | | | | | |
|---|---|---|---|---|---|---|---|---|---|---|---|---|
| **treesSherman** | Condit 1998 | Trees | Chagres river, Panama | 0.06 | Nowember 18, 1997 - January 23, 1998 | 66 | Trees observed from the plot on a given day | 32 | 24441 | 238 | 103 | 13.87 |
| **treesTapajos** | Goncalves et al. 2018 | Trees | Tapajos National Forest, Para, Brazil | - | August 31, 2010 - September 16, 2010 | 16 | Trees inside a plot | 30 | 3371 | 140 | 24 | 17.14 |

### References


Barea-Azcón, J.M. (2021) Dataset of butterfly monitoring in Sierra Nevada (Spain). Version 1.1. Sierra Nevada Global Change Observatory.Andalusian Environmental Center, University of Granada, Regional Government of Andalusia. Sampling event dataset https://doi.org/10.15470/tc3gdq accessed via GBIF.org on 2021-09-10.

Bateman, H., Childers, D., Warren, P. (2018) Point-count bird censusing: long-term monitoring of bird abundance and diversity along the Salt River in the greater Phoenixmetropolitan area, ongoing since 2013 (Reformatted to ecocomDP Design Pattern) ver 1. Environmental Data Initiative. https://doi.org/10.6073/pasta/23a69b9940326530defdf1831f467a24 (Accessed 2020-12-11).

Baturina, M., Kaygorodova, I., Makarevich, O. (2020) The fauna of annelid worms (Oligochaeta, Hirudinea) of the Naroch lakes system (Belarus). Institute of Biology of Komi Scientific Centre of the Ural Branch of the Russian Academy of Sciences. Sampling event dataset https://doi.org/10.15468/4ajykm accessed via GBIF.org on 2021-09-10.

Beja, P., Motta-Ferreira, M., Filpe, A.F., Carona, S., Henrique, P., Severino, R., Ivone, S., Henrique, S., Prata, D., Lopes, J., Guilherme, J., Barradas, J., Cereja, R., Pace, G., Rosa, I., Pinto, M., Agudelo, W., Buzzo, G., Sá, F., Quaglietta, L., Pereira, N., Múrias, T. (2021) LTER Baixo Sabor: Long term monitoring of freshwater fish - Sabor watershed [2012 - 2020]. Version 1.3. CIBIO (Research Center in Biodiversity and Genetic Resources) Portugal. Sampling event dataset https://doi.org/10.15468/pep8ma accessed via GBIF.org on 2021-10-07.

Berrios, Hazel (2020) Species richness and abundance of vascular epiphytes along an elevation gradient, Dryad, Dataset, https://doi.org/10.5061/dryad.bzkh1896h

Bitušík, P., Novikmec, M., Hamerlik, L. (2020) Chironomids (Insecta, Diptera, Chironomidae) from alpine lakes in the Eastern Carpathians with comments on newly-recorded species from Ukraine. Biodiversity Data Journal 8: e49378. https://doi.org/10.3897/BDJ.8.e49378

Brooks, A., (2019) Moorea Coral Reef MCR LTER: Coral Reef: Long-term Population and Community Dynamics: Fishes, ongoing since 2005 ver 57. Environmental Data Initiative. https://doi.org/10.6073/pasta/ef2b0709cb84eb37a33a9895de5f6f7e (Accessed 2021-01-08).

Cardoso, P., Gaspar, C., Pereira, L.C., Silva, I., Henriques, S.S., Silva, R.R. & Sousa, P. (2008a) Assessing spider species richness and composition in Mediterranean cork oak forests. Acta Oecologica, 33: 114-127. https://doi.org/10.1016/j.actao.2007.10.003

Cardoso, P., Scharff, N., Gaspar, C., Henriques, S.S., Carvalho, R., Castro, P.H., Schmidt, J.B., Silva, I., Szuts, T., Castro, A. & Crespo, L.C. (2008b) Rapid biodiversity assessment of spiders (Araneae) using semi-quantitative sampling: a case study in a Mediterranean forest. Insect Conservation and Diversity, 1: 71-84. https://doi.org/10.1111/j.1752-4598.2007.00008.x

Chatzinikolaou, E., Arvanitidis, C., (2017). Benthic communities and environmental parameters in three Mediterranean ports (Sardinia, Crete, Tunisia). https://doi.org/10.15468/xrlqx4 accessed via GBIF.org on 2021-05-04.

Choi, H-A., Seliger, B., Moores, N., Borzée, A., Yoon, CHK. (2020) Avian Surveys in the Korean Inner Border Area, Gimpo, Republic of Korea. Biodiversity Data Journal 8: e56219. https://doi.org/10.3897/BDJ.8.e56219

Condit, R. (1998) Ecological implications of changes in drought patterns: shifts in forest composition in Panama. Climatic Change, 39: 413-427. https://doi.org/10.1023/A:1005395806800

Cornthwaite, M. (2021) DFO Pacific Inside South Hard Bottom Longline Surveys. Version 1.1. Fisheries and Oceans Canada. Sampling event dataset https://doi.org/10.15468/5d545y accessed via GBIF.org on 2021-09-24.

Costa, H., Salgueiro, N., Marques, A. T., Coelho, H., Gonçalves, R., Brás, L., Monteiro, B., Ferreira, C., Paula, J., Oliveira, J., Puga, J., Silva, M., Neves, T., Pereira, F., Magalhães, J., Romão, F., Rosa, L., Sousa, D., Pereira, P., Cabral, J. A., Vale-Gonçalves, H. M., Barros, P., Múrias, T. (2018) EDP Foz Tua: Amphibians - Ecological Monitoring Program [2010-2017]. EDP - Energias de Portugal. Sampling event dataset https://doi.org/10.15468/uo5uyu accessed via GBIF.org on 2021-10-07.

Costa, H., Salgueiro, N., Marques, A. T., Coelho, H., Monteiro, B., Brás, L., Gonçalves, R., Oliveira, J., Silva, M., Sousa, D., Caetano, M., Romão, F., Santos, J., Peixoto, M., Múrias, T. (2020) EDP Foz-Tua: Dragonflies and Damselflies (Odonata) - Ecological Monitoring

Program (2011-2017). Version 1.1. EDP - Energias de Portugal. Sampling event dataset https://doi.org/10.15468/ywwyjd accessed via GBIF.org on 2021-05-11.

Coulibaly, D., Stein, K. (2018) Managing West African Bees in the implementation of a first reference collection: Bees caught in three areas of Burkina Faso. Station d'Ecologie de Lamto. Sampling event dataset https://doi.org/10.15468/njbmsp accessed via GBIF.org on 2021-10-07.

Dekoninck, W., Versteirt, V., Van Bortel, W., Brosens, D. (2021) MODIRISK: Monitoring of Mosquito Vectors, Longitudinal study. Version 1.10. Royal Belgian Institute of Natural Sciences. Sampling event dataset https://doi.org/10.15468/rwsozv accessed via GBIF.org on 2021-07-19.

El Zein, H. (2020) Vascular plants of the Valleys of Hell, Danniye-Akkar, Lebanon, 2019. Version 1.7. Wadi ez-Zouhour. Sampling event dataset https://doi.org/10.15468/jh6usj accessed via GBIF.org on 2021-09-10.

Ellison, A., Gotelli, N. (2018) Inventory of Ants at the Black Rock Forest in Cornwall NY since 2006 ver 23. Environmental Data Initiative. https://doi.org/10.6073/pasta/f9eb0e3bcc9a32b8e87c64ba9a5bf4bc (Accessed 2021-01-04).

Gastauer, M. (2016) corrego_fazendinha-gallery-forest_v2. Laboratory of Ecology and Evolution of Plants, at Universidade Federal de Vicosa. Occurrence dataset https://doi.org/10.15468/judbja accessed via GBIF.org on 2021-10-07

Girotto, A., Groom, Q. (2018) Bird Monitoring 2017, Meise Botanic Garden. Version 1.6. Meise Botanic Garden. Sampling event dataset https://doi.org/10.15468/xecp6u accessed via GBIF.org on 2021-09-24.

Goncalves, F.G., Treuhaft, R.N., Dos santos, J.R., Graca, P., Almeida A., Law, B.E. (2018) Tree Inventory and Biometry Measurements, Tapajos National Forest, Para, Brazil, 2010. ORNL DAAC, Oak Ridge, Tennessee, USA. https://doi.org/10.3334/ORNLDAAC/1552,

Goulart, S., Barreiros, J.P., Brito, M.R., Santos, S., Pimentel, C., Nogueira, E.C., Borges, P.A.V. (2019) Birds from Praia da Vitória marshes (Terceira, Azores, Portugal). Universidade dos Açores. Sampling event dataset https://doi.org/10.15468/dnffus accessed via GBIF.org on 2021-01-05.

Haarstad, J. (2018) Orthoptera species abundance: Trophic Structure: Insect Species Diversity, Abundance and Body Size ver 8. Environmental Data Initiative. https://doi.org/10.6073/pasta/11223e0d49d878ddd211f934ddef86c7 (Accessed 2021-02-17).

Herrera Varon, Y (2020) Mamíferos y aves medianos y grandes de los Montes de María - Programa Riqueza Natural (USAID). Instituto de Investigación de Recursos Biológicos Alexander von Humboldt. Sampling event dataset https://doi.org/10.15472/vhddix accessed via GBIF.org on 2021-06-01.

Kano, Y., Musikasinthorn, P., Iwata, A., Tun, S., Yun, L., Win, S., Matsui, S., Tabata, R., Yamasaki, T., Watanabe, K. (2016) A dataset of fishes in and around Inle Lake, an ancient lake of Myanmar, with DNA barcoding, photo images and CT/3D models. Biodiversity Data Journal 4: e10539. https://doi.org/10.3897/BDJ.4.e10539

Kascelan, S., Mandic, S., Radovic I., Krpo-Cetkovic, J. (2015) Institute of Marine Biology - University of Montenegro; University of Belgrade. Echinodermata of Montenegro (South Adriatic). https://doi.org/10.14284/30 accessed via GBIF.org on 2021-06-02.

Langeraert, W., Brosens, D. (2020) Land and freshwater molluscs of Gran Canaria (Spain). Version 1.13. Ghent University. Occurrence dataset https://doi.org/10.15468/ny1f9n accessed via GBIF.org on 2021-05-19.

Lind, L., Nilsson, C. (2015) Data from: Vegetation patterns in small boreal streams relate to ice and winter floods, Dryad, Dataset, https://doi.org/10.5061/dryad.pn63g

Makarov, K., Matalin, A. (2020) Carabid beetles of the environs of the Elton Lake: fauna, population dynamics, demography. Version 1.2. Moscow Pedagogical State University (MPGU). Sampling event dataset https://doi.org/10.15468/a8weeh accessed via GBIF.org on 2021-05-11.

Melo, C.D., Walker, C., Freitas, H., Machado, A.C, Borges, P.A.V. (2020) Distribution of arbuscular mycorrhizal fungi (AMF) in Terceira and São Miguel Islands (Azores). Biodiversity Data Journal 8: e49759. https://doi.org/10.3897/BDJ.8.e49759

Mensah, S., Salako, V., Seifert, T. (2020) Data from: Structural complexity and large-sized trees explain shifting species richness and carbon relationship across vegetation types, Dryad, Dataset, https://doi.org/10.5061/dryad.pvmcvdnhr

Mosie, I., Makati, K. (2020) Long Term Time-Series Data on Fish Monitoring by Okavango Research Institute, Botswana. Version 1.3. Okavango Research Institute. Sampling event dataset https://doi.org/10.15468/4vvwzc accessed via GBIF.org on 2021-09-10.

Noel, G., Bonnet, J., Everaerts, S., Danel, A., Calderan, A., de Liedekerke, A., de Montpellier d'Annevoie, C., Francis, F., Serteyn, L. (2021) Distribution of wild bee (Hymenoptera: Anthophila) and hoverfly (Diptera: Syrphidae) communities within farms undergoing ecological transition. Biodiversity Data Journal 9: e60665. https://doi.org/10.3897/BDJ.9.e60665

Nyirambangutse, B., et al. (2017) Data from: Carbon stocks and dynamics at different successional stages in an Afromontane tropical forest, Dryad, Dataset, https://doi.org/10.5061/dryad.b5b4h

Parker, P.D.M. (2019) FBIP: Insectivorous bat monitoring in the Kruger National Park. South African National Biodiversity Institute. Occurrence dataset https://doi.org/10.15468/ff5mzy accessed via GBIF.org on 2021-05-03

Paz-Ríos, C.E., Sosa-López, A., Torres-Rojas, Y.E., Ramos-Miranda, J., del Río-Rodríguez, R.E. (2021) Fish species richness in the Terminos Lagoon: An occurrence data compilation of four sampling campaigns along a multidecadal series. Biodiversity Data Journal 9: e65317. https://doi.org/10.3897/BDJ.9.e65317

Pina, S. (2017) The Orthoptera of Castro Verde Special Protection Area (Southern Portugal). CIBIO (Research Center in Biodiversity and Genetic Resources) Portugal. Sampling event dataset https://doi.org/10.15468/byd0kt accessed via GBIF.org on 2021-10-03.

Pyke, C.R., Condit, R., Aguilar, S., Lao, S. (2001) Floristic composition across a climatic gradient in a neotropical lowland forest. Journal of Vegetation Science, 12: 553-566. https://doi-org.libproxy.helsinki.fi/10.2307/3237007

Ratti (2004) Master Thesis / Fraser Valley, British Columbia. York University. Version 1.0. Occurrence dataset https://doi.org/10.15468/c9bdau accessed via GBIF.org on 2021-10-03.

Ross, D.N. (2014) Andrews Forest LTER Site. Spatial and temporal distribution and abundance of butterflies in the Andrews Experimental Forest, 1994-1996 ver 5. Environmental Data Initiative. https://doi.org/10.6073/pasta/b75017cddc991a02d3ed3302578b5b45 (Accessed 2020-12-09).

Seastedt, T. (2019) Krummholz island plant species density data for East of Tvan, 2000. ver 3. Environmental Data Initiative. https://doi.org/10.6073/pasta/be46adf5b277b2be25639e0882be0898 (Accessed 2021-01-06).

Schrader, J., Moeljono, S., Tambing, J., Sattler, C., Kreft, H. (2020) A new dataset on plant occurrences on small islands, including species abundances and functional traits across different spatial scales. Biodiversity Data Journal 8: e55275. https://doi.org/10.3897/BDJ.8.e55275

Trubina, M., Nesterkov, A. (2021) Diversity and distribution of vascular plants within the treeline ecotone in Mount Iremel (Southern Urals, Russia). Version 1.4. Institute of Plant and Animal Ecology (IPAE). Sampling event dataset https://doi.org/10.15468/6hsht5 accessed via GBIF.org on 2021-05-06.

Vega-Álvarez, J., García-Rodríguez, J.A., Cayuela, L. (2019) Facilitation beyond species richness. J. Ecol. 107: 722– 734. https://doi.org/10.1111/1365-2745.13072

Zhang, K., Lin, S., Ji, Y., Yang, C., Wang, X., Yang, C., Wang, H., Jiang, H., Harrison, R.D., Yu, D.W. (2016) Plant diversity accurately predicts insect diversity in two tropical landscapes. Mol Ecol, 25: 4407-4419. https://doi-org.libproxy.helsinki.fi/10.1111/mec.13770

Zimmerman, J.K., Liza, S.C., Thompson J., Uriarte, M., Brokaw, N. (2010) Patch dynamics and community metastability of a subtropical forest: compound effects of natural disturbance and human land use. Landscape Ecology 25: 1099-1111. https://doi.org/10.1007/s10980-010-9486-x
