## Appendix B for "Accuracy of non-parametric species richness estimators across taxa and regions"

**Table B1.** Species richness estimates for each dataset and the respective mean.

| Dataset | Jack1ab | Jack1abP | Jack1in | Jack1inP | Jack2ab | jack2abP | Jack2in | Jack2inP | Chao1 | Chao1P | Chao2 | Chao2P | Mean |
|---|---|---|---|---|---|---|---|---|---|---|---|---|---|
| amphibiansTua | 13.00 | 13.09 | 13.00 | 13.09 | 14.00 | 14.10 | 13.99 | 14.09 | 12.00 | 12.08 | 12.00 | 12.08 | 13.04 |
| annelidaBelarus | 63.00 | 65.82 | 69.88 | 78.26 | 69.00 | 72.09 | 80.79 | 90.47 | 61.17 | 63.90 | 71.13 | 79.65 | 72.10 |
| batsKruger | 24.00 | 24.05 | 24.86 | 25.05 | 25.00 | 25.05 | 26.57 | 26.77 | 23.00 | 23.04 | 24.00 | 24.18 | 24.63 |
| beesCanada | 71.00 | 72.14 | 72.89 | 74.73 | 71.00 | 72.14 | 76.87 | 78.80 | 66.11 | 67.18 | 69.43 | 71.18 | 71.96 |
| beesBurkina1 | 36.00 | 37.98 | 38.10 | 36.00 | 36.00 | 35.08 | 35.19 | 36.00 | 36.00 | 36.17 | 36.28 | 36.23 | 36.25 |
| beesBurkina2 | 38.03 | 45.91 | 48.63 | 39.00 | 39.03 | 52.79 | 55.92 | 37.00 | 37.03 | 49.00 | 51.90 | 44.35 | 44.88 |
| beesBurkina3 | 37.00 | 42.93 | 44.06 | 35.00 | 35.00 | 43.97 | 45.12 | 37.00 | 37.00 | 39.50 | 40.54 | 39.51 | 39.72 |
| beetlesRussia | 168.00 | 171.07 | 170.89 | 175.02 | 172.00 | 175.14 | 176.91 | 181.19 | 159.18 | 162.08 | 162.06 | 165.97 | 169.96 |
| benthicMed1 | 204.00 | 211.06 | 216.98 | 232.50 | 214.00 | 221.41 | 235.72 | 252.58 | 193.57 | 200.27 | 208.96 | 223.91 | 217.91 |
| benthicMed2 | 161.00 | 169.44 | 171.02 | 187.78 | 174.00 | 183.13 | 189.63 | 208.21 | 155.17 | 163.30 | 166.65 | 182.98 | 176.03 |
| benthicMed3 | 165.00 | 170.26 | 174.53 | 185.44 | 172.00 | 177.48 | 189.40 | 201.23 | 155.79 | 160.76 | 168.33 | 178.85 | 174.92 |
| birdsAzores | 118.00 | 119.25 | 117.99 | 119.24 | 123.00 | 124.30 | 121.00 | 122.27 | 114.86 | 116.07 | 113.11 | 114.31 | 118.62 |
| birdsBelgium | 40.00 | 40.26 | 40.89 | 41.37 | 38.00 | 38.25 | 41.00 | 41.48 | 37.50 | 37.75 | 38.20 | 38.65 | 39.44 |
| birdsKorea | 127.00 | 129.28 | 147.05 | 163.10 | 133.00 | 135.39 | 162.41 | 180.14 | 122.50 | 124.70 | 142.27 | 157.80 | 143.72 |
| chiroUkraine | 40.00 | 40.82 | 49.88 | 61.64 | 43.00 | 43.88 | 56.84 | 70.25 | 38.33 | 39.12 | 48.60 | 60.07 | 49.37 |
| culicidaeBelgium | 19.00 | 19.26 | 19.89 | 20.51 | 20.00 | 20.28 | 22.68 | 23.38 | 17.50 | 17.74 | 20.00 | 20.62 | 20.07 |
| echinoMontenegro | 52.00 | 52.08 | 63.61 | 68.60 | 51.00 | 51.08 | 71.33 | 76.92 | 50.25 | 50.33 | 63.00 | 67.94 | 59.85 |
| epiphytesNicaragua | 84.00 | 84.21 | 116.08 | 140.90 | 85.00 | 85.21 | 141.97 | 172.34 | 81.50 | 81.70 | 140.55 | 170.61 | 115.34 |
| fishBotswana1 | 54.00 | 54.35 | 55.94 | 56.74 | 58.00 | 58.37 | 58.91 | 59.76 | 56.00 | 56.36 | 53.75 | 54.52 | 56.39 |
| fishBotswana2 | 29.00 | 29.39 | 29.83 | 30.53 | 32.00 | 32.43 | 31.74 | 32.49 | 29.00 | 29.39 | 28.00 | 28.66 | 30.20 |

| | | | | | | | | | | | | | |
|---|---|---|---|---|---|---|---|---|---|---|---|---|---|
| **fishBotswana3** | 54.00 | 54.35 | 55.94 | 56.74 | 58.00 | 58.37 | 58.91 | 59.76 | 56.00 | 56.36 | 53.75 | 54.52 | 56.39 |
| **fishCanada** | 40.00 | 40.26 | 40.89 | 41.37 | 38.00 | 38.25 | 41.00 | 41.48 | 37.50 | 37.75 | 38.20 | 38.65 | 39.44 |
| **fishMexico** | 167.00 | 172.68 | 176.84 | 189.02 | 187.00 | 193.36 | 199.74 | 213.49 | 187.43 | 193.80 | 188.57 | 201.56 | 189.21 |
| **fishMyanmar** | 43.00 | 43.10 | 44.91 | 45.34 | 44.00 | 44.10 | 41.27 | 41.66 | 41.50 | 41.60 | 41.67 | 42.06 | 42.85 |
| **fishSabor** | 15.00 | 15.08 | 15.00 | 15.07 | 16.00 | 16.08 | 15.00 | 15.08 | 14.00 | 14.07 | 14.00 | 14.07 | 14.87 |
| **fungiAzores** | 38.00 | 38.03 | 41.98 | 42.74 | 39.00 | 39.03 | 42.99 | 43.77 | 37.00 | 37.03 | 39.00 | 39.71 | 39.86 |
| **lepidopteraSpain** | 103.00 | 103.01 | 104.00 | 104.04 | 103.00 | 103.01 | 106.00 | 106.04 | 102.00 | 102.01 | 103.00 | 103.04 | 103.51 |
| **LTER_ants** | 48.00 | 49.40 | 51.80 | 55.52 | 51.00 | 52.49 | 58.61 | 62.83 | 45.20 | 46.52 | 52.00 | 55.74 | 52.43 |
| **LTER_birds** | 207.00 | 209.37 | 223.80 | 232.57 | 207.00 | 209.37 | 248.60 | 258.34 | 196.05 | 198.29 | 238.23 | 247.56 | 223.01 |
| **LTER_butterflies** | 77.00 | 78.35 | 77.88 | 79.57 | 83.00 | 84.45 | 83.79 | 85.60 | 77.00 | 78.35 | 77.00 | 78.67 | 80.05 |
| **LTER_Orthoptera** | 60.00 | 61.42 | 62.78 | 65.58 | 66.00 | 67.56 | 70.51 | 73.66 | 61.33 | 62.79 | 65.75 | 68.69 | 65.51 |
| **LTER_plants** | 48.00 | 48.65 | 49.50 | 50.81 | 51.00 | 51.69 | 50.75 | 52.10 | 46.33 | 46.96 | 46.00 | 47.22 | 49.08 |
| **LTER_reefFish1** | 303.00 | 308.86 | 313.25 | 323.88 | 320.00 | 326.19 | 334.61 | 345.96 | 297.71 | 303.47 | 308.00 | 318.45 | 316.95 |
| **LTER_reefFish2** | 295.00 | 300.35 | 305.32 | 315.30 | 312.00 | 317.65 | 325.72 | 336.36 | 291.32 | 296.59 | 300.04 | 309.84 | 308.79 |
| **LTER_reefFish3** | 294.00 | 300.09 | 311.07 | 325.84 | 318.00 | 324.59 | 347.05 | 363.52 | 304.57 | 310.88 | 338.05 | 354.10 | 324.31 |
| **LTER_reefFish4** | 289.00 | 293.80 | 304.08 | 315.68 | 309.00 | 314.13 | 328.96 | 341.51 | 293.71 | 298.59 | 305.00 | 316.63 | 309.18 |
| **LTER_reefFish5** | 281.00 | 285.01 | 296.38 | 306.77 | 296.00 | 300.23 | 321.28 | 332.55 | 278.19 | 282.16 | 300.14 | 310.66 | 299.20 |
| **LTER_reefFish6** | 280.00 | 284.03 | 290.44 | 298.64 | 300.00 | 304.32 | 307.97 | 316.66 | 289.55 | 293.71 | 284.44 | 292.47 | 295.19 |
| **mammalsColombia** | 26.00 | 26.04 | 25.98 | 26.03 | 25.00 | 25.04 | 25.05 | 25.09 | 25.00 | 25.04 | 25.00 | 25.04 | 25.36 |
| **molluscsCanaries** | 56.00 | 56.08 | 69.54 | 75.65 | 55.00 | 55.08 | 77.31 | 84.09 | 54.25 | 54.32 | 67.33 | 73.24 | 64.82 |
| **odonataTua** | 44.00 | 44.44 | 44.98 | 45.68 | 44.00 | 44.44 | 44.01 | 44.70 | 41.20 | 41.61 | 41.43 | 42.08 | 43.55 |
| **orthopteraCastroVerde** | 39.00 | 39.51 | 39.98 | 40.80 | 38.00 | 38.50 | 38.02 | 38.79 | 36.00 | 36.47 | 36.25 | 36.99 | 38.19 |
| **plantsIndonesia** | 61.00 | 61.30 | 66.75 | 68.80 | 53.00 | 53.26 | 62.37 | 64.28 | 57.46 | 57.74 | 59.81 | 61.65 | 60.62 |
| **plantsLebanon** | 479.00 | 480.01 | 556.38 | 582.38 | 470.00 | 470.99 | 599.19 | 627.19 | 464.77 | 465.75 | 543.11 | 568.48 | 525.60 |

| | | | | | | | | | | | | | |
|---|---|---|---|---|---|---|---|---|---|---|---|---|---|
| **plantsRussia** | 82.00 | 83.25 | 81.99 | 83.23 | 87.00 | 88.32 | 86.98 | 88.30 | 80.20 | 81.42 | 80.20 | 81.42 | 83.69 |
| **plantsSpain** | 156.00 | 156.73 | 158.98 | 160.24 | 155.00 | 155.73 | 160.99 | 162.27 | 149.75 | 150.45 | 152.50 | 153.71 | 156.03 |
| **plantsSweden1** | 100.00 | 102.65 | 105.90 | 111.62 | 109.00 | 111.89 | 111.91 | 117.96 | 101.17 | 103.85 | 98.67 | 104.00 | 106.55 |
| **plantsSweden2** | 85.00 | 85.21 | 92.94 | 94.98 | 84.00 | 84.20 | 94.97 | 97.05 | 82.00 | 82.20 | 87.00 | 88.91 | 88.20 |
| **plantsSweden3** | 98.00 | 98.42 | 109.89 | 114.10 | 98.00 | 98.42 | 116.87 | 121.35 | 94.14 | 94.54 | 104.75 | 108.76 | 104.77 |
| **polinatorsBelgium** | 145.00 | 151.29 | 150.22 | 162.28 | 154.00 | 160.68 | 163.06 | 176.15 | 137.65 | 143.62 | 146.71 | 158.49 | 154.10 |
| **spidersArrabida** | 176.00 | 181.29 | 178.91 | 178.91 | 177.00 | 182.32 | 183.95 | 190.83 | 162.50 | 167.38 | 166.24 | 172.45 | 176.48 |
| **spidersGeres** | 223.00 | 231.82 | 225.88 | 236.32 | 235.00 | 244.30 | 240.86 | 252.00 | 211.62 | 219.99 | 216.00 | 225.99 | 230.23 |
| **treesBCI** | 244.00 | 245.74 | 245.58 | 247.72 | 250.00 | 251.78 | 247.87 | 250.03 | 237.21 | 238.91 | 235.50 | 237.55 | 244.32 |
| **treesBenin** | 48.00 | 48.09 | 47.95 | 48.04 | 48.00 | 48.09 | 47.07 | 47.16 | 46.33 | 46.42 | 46.25 | 46.34 | 47.31 |
| **treesBrazil** | 95.00 | 98.34 | 96.67 | 101.03 | 102.00 | 105.59 | 104.53 | 109.25 | 91.67 | 94.89 | 93.60 | 97.83 | 99.20 |
| **treesChina1** | 941.00 | 966.94 | 1153.18 | 1381.39 | 1024.00 | 1052.23 | 1355.09 | 1623.26 | 978.37 | 1005.34 | 1241.20 | 1486.83 | 1184.07 |
| **treesChina2** | 427.00 | 436.73 | 482.03 | 528.35 | 444.00 | 454.12 | 524.46 | 574.86 | 409.50 | 418.83 | 460.79 | 505.07 | 472.15 |
| **treesCocoli** | 205.00 | 211.02 | 209.80 | 218.68 | 214.00 | 220.29 | 224.48 | 233.98 | 194.77 | 200.50 | 203.64 | 212.25 | 212.37 |
| **treesPuertoRico** | 160.00 | 161.47 | 164.95 | 167.75 | 165.00 | 166.52 | 174.92 | 177.89 | 155.10 | 156.53 | 163.10 | 165.86 | 164.92 |
| **treesRwanda** | 94.00 | 94.81 | 108.40 | 116.84 | 95.00 | 95.82 | 114.71 | 123.65 | 89.50 | 90.27 | 100.53 | 108.36 | 102.66 |
| **treesSherman** | 271.00 | 276.21 | 271.91 | 277.79 | 288.00 | 293.54 | 289.30 | 295.55 | 269.06 | 274.23 | 271.06 | 276.92 | 279.55 |
| **treesTapajos** | 164.00 | 168.82 | 166.10 | 172.28 | 172.00 | 177.05 | 173.28 | 179.72 | 156.24 | 160.83 | 156.71 | 162.54 | 167.46 |
